# Substrate deformations induce directed keratinocyte migration

**DOI:** 10.1101/249037

**Authors:** Hoda Zarkoob, Sathivel Chinnathambi, John C. Selby, Edward A. Sander

## Abstract

Cell migration is an essential part of many (patho)physiological processes in the body, including keratinocyte re-epithelialization of healing wounds. Recent interest in the mechanobiology of tissues suggests that physical forces and mechanical cues from the wound bed (in addition to biochemical signals) may also play an important role in the healing process. Previously, we explored this possibility and found that polyacrylamide (PA) gel stiffness affected primary human keratinocyte behavior and that mechanical deformations in soft (~1.2 kPA) PA gels produced by neighboring cells appeared to influence the process of *de novo* epithelial sheet formation. In order to clearly demonstrate that keratinocytes do respond to such deformations, we conducted a series of experiments where we observed the response of single keratinocytes to a prescribed local substrate deformation that mimicked a neighboring cell or evolving multicellular aggregate via a servo-controlled microneedle. We also examined the effect of adding either Y27632, a rho kinase inhibitor, or blebbistatin, a non-muscle myosin II inhibitor, on the response of the cells to PA gel deformations. The results of this study indicate that keratinocytes do sense and respond to mechanical signals comparable to those that originate from substrate displacements imposed by neighboring cells, a finding that could have important implications for the process of keratinocyte re-epithelialization that takes place during normal and pathologic wound healing. Furthermore, the Rho/ROCK pathway and the engagement of NM II are both essential to the observed process of substrate deformation-directed keratinocyte migration.

## INTRODUCTION

Cell migration is an essential part of many physiological and pathological processes in the body, including tissue morphogenesis [1], inflammation [2], and wound healing [3]. Re-epithelialization during cutaneous wound healing is a result of migration, proliferation, and differentiation of the peripheral keratinocytes along the edge of the wound [4-6]. Aberrant or delayed re-epithelialization increases the risk of infection and the severity of scar tissue formation, and it contributes to the development and persistence of chronic wounds [4, 7, 8]. The regulation of re-epithelialization and keratinocyte migration is understood largely in terms of biochemical signals [6]. Recent interest in the mechanobiology of tissues, however, suggests that physical forces and mechanical cues from the wound bed could also play an important role in the healing process [4, 9-11].

The biophysical mechanisms by which cells sense and respond to mechanical signals, however, remain poorly understood. In addition to the multi-protein complexes that form at the anchoring junctions between the cell and the substrate [12, 13], actin-myosin interactions with the cytoskeleton remain a focal point of study for elucidating the mechanisms involved in mechanosensing, particularly those involving non-muscle myosin II (NM II) [14-16]. NM II engages with actin to generate cytoskeletal traction forces that are transmitted through an integrated complex of focal adhesion associated proteins to the extracellular matrix (ECM) via integrins, the main transmembrane receptors for ECM proteins [17, 18]. An increase in ECM/substrate stiffness recruits more integrin-associated structural and signaling proteins to the site of adhesion, including focal adhesion kinase (FAK). Increased FAK activity at these adhesive contacts also stimulates the Rho/ROCK (Rho kinase) pathway. Rho is a family of small GTPases that activate ROCK, which in turn regulates myosin light chain (MLC) phosphorylation, NM II engagement, and contraction with actin, which leads to transmembrane force generation that promotes cell migration [19, 20] or substrate displacement [21, 22]. Collectively, cellular components that contribute to myosin contraction, such as Rho/ROCK, appear to be vital to mechanosensing for a range of cell phenotypes [20, 22], possibly through a mechanism involving substrate deformation dependent mechanical feedback. Indeed, several studies have indicated that Rho/ROCK and NMII are important to keratinocyte behavior, including regulation of keratinocyte differentiation [23-28].

It is important to understand not only the basic mechanisms underlying cellular mechanosensation, but also how this process controls individual and coordinated physiological and pathophysiological cellular behaviors, such as those involved in tissue self-assembly [29-32], ECM remodeling [33, 34], wound healing [4, 9-11], fibrosis [35, 36], and cancer metastasis [37-41]. For example, the influence of ECM remodeling and stiffness on tumor progression has been the subject of some recent cancer studies [39, 41]. With respect to wound healing, Zarkoob *et al*. explored previously how substrate stiffness affected primary human keratinocyte behavior and found that mechanical cues influenced the process of *de novo* epithelial sheet formation [42]. Specifically, the process by which human keratinocyte formed multicellular aggregates was significantly different on soft (~ 1.2 kPa) versus stiff (~24 kPa) polyacrylamide (PA) gels [42]. Keratinocytes on soft PA gels exhibited smaller spread contact areas, increased migration velocities, increased rates of aggregate formation, and more cells per aggregate, respectively [42]. In addition, the keratinocytes on soft PA gels appeared to migrate directly towards an evolving multicellular aggregate in a cooperative manner, ostensibly in response to mechanical cues that propagated through the deforming substrate from the aggregate. Such directed migration was not apparent on the stiff PA gels.

These observations were near completely analogous to those published by Reinhart-King *et al*. with respect to bovine aortic endothelial cells plated at low density on 2.5 – 5.5 kPa PA gels [43]. Both Reinhart-King *et al*. and Zarkoob *et al*. [42] hypothesized that such enhanced cooperativity is due to the ability of a cell to sense and respond to substrate deformations induced by the neighboring cells. In such a scenario, because each cell likely modulates the tractions it exerts on the substrate in response to those of the adjacent contracting cell, the mechanical environment is spatially heterogeneous and temporally dynamic, which makes it difficult to correlate cell behavior unambiguously to mechanical signals communicated through the substrate from neighboring cells.

In order to clearly demonstrate that keratinocytes do respond to such deformations, we conducted a series of experiments where we observed the response of single keratinocytes to a prescribed local substrate deformation that mimicked a neighboring cell or evolving multicellular aggregate via a servo-controlled microneedle. We also examined the effect of adding either Y27632, a rho kinase inhibitor, or blebbistatin, a NM II inhibitor, on the response of the cells to PA gel deformations. Both of these chemicals are commonly used to investigate the role that internal force generation plays in the mechanosensing process [44-47]. The results of this study indicate that keratinocytes do sense and respond to mechanical signals comparable to those that originate from substrate displacements imposed by neighboring cells, a finding that could have important implications for the process of keratinocyte re-epithelialization that takes place during normal and pathologic wound healing.

## MATERIAL AND METHODS

### Polyacrylamide Gel Preparation

Thin polyacrylamide (PA) gels approximately 100 μm in thickness were polymerized on the surfaces of glass bottom Petri dishes (MatTec Corp., Ashland, MA) as described previously [42, 48]. Briefly, the ratios of 2.0%/0.25% and 7.5%/0.3% acrylamide/bis-acrylamide were used to fabricate soft (~1.2 kPa) and stiff (~24 kPa) PA gels, respectively (nominal stiffness based on gel formulations published elsewhere [49, 50]). FluoroSpheres® Carboxylate-Modified Microspheres (#F8812, Life Technologies, Carlsbad, CA) measuring 0.5 μm in diameter were embedded inside the gels to facilitate deformation tracking. Pepsin-digested collagen was attached covalently to the surface of the gel with Sulfo-SANPAH. The samples were sterilized under UV light for 15 minutes and stored at 4 °C for later use.

### Cell Culture

Neonatal human epidermal keratinocytes (HEKn) (Fisher Scientific, Waltham, MA) were cultured in keratinocyte serum free medium (KSFM) (Invitrogen) supplemented with 1% penicillin-streptomycin and 0.1% amphotericin B in a humidified incubator maintained at 37 °C and 5%/95% CO_2_/air. Passage three cells were plated at a low density of approximately 200 cells/cm^2^ onto the surface of the PA gels. The calcium of the medium was then elevated from a baseline concentration of ~0.09 mM to ~1.2 mM by adding CaCl_2_ to the medium in order to trigger cell-cell anchoring junction assembly and de novo epithelial sheet formation via the calcium switch [42, 51, 52]. HEKn were allowed to attach and equilibrate for three hours in the incubator before imaging with the microscope.

### Time-Lapse Live Cell Imaging

Once the samples had equilibrated, they were transferred to a temperature-controlled microscope enclosure (CO_2_ Microscope Cage Incubation System, Okolab, Pozzuoli, Italy) with recirculating air that integrates with a micro-environmental gas chamber (H201-K-Frame, Okolab, Pozzuoli, Italy) (Fig. 1). This setup provides 37 °C, humidified air with 5% CO_2_ via a manual gas mixer and pump to the samples so that extended time-lapse imaging is possible. Time-lapse images were acquired with a Nikon Ti-E inverted microscope equipped with wide-field epifluorescence and differential interference contrast (DIC) microscopy capabilities and a DS-Qi1 Nikon camera. DIC and fluorescent images pairs were acquired with a CFI plan Apo 10X DIC objective combined with a 1.5X magnifier.

**Figure 1.**
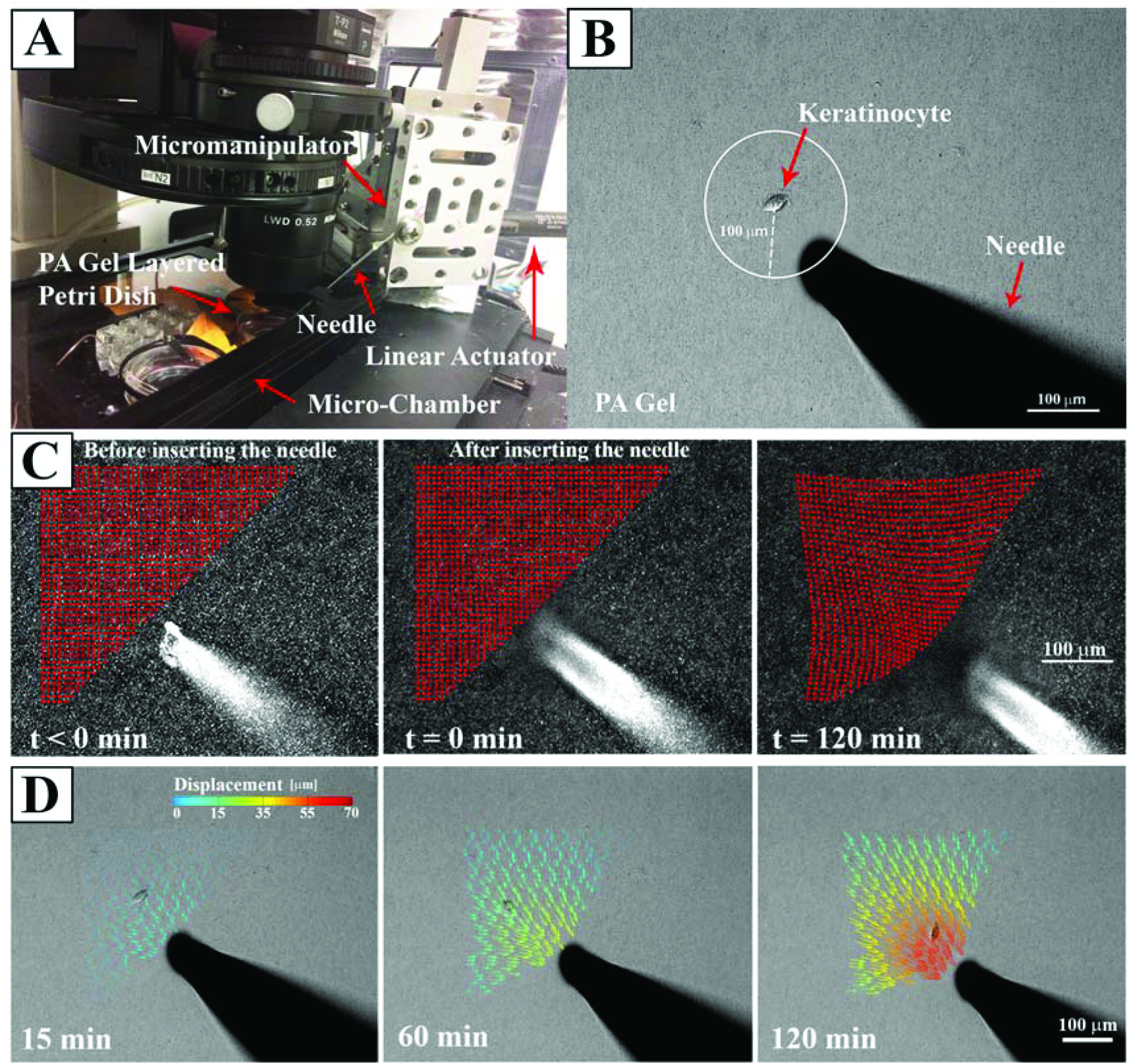
Experimental setup to apply controlled mechanical deformations to PA gels. (A) The PA gel is placed within a microscope-mounted micro-chamber contained with an environmentally controlled enclosure. A servo-controlled liner actuator drives needle movement via a high-precision micromanipulator. (B) A single keratinocyte is identified and the needle is inserted a prescribed distance away from the centroid of the cell, in this instance ~100 μm. (C) Substrate deformation in response to needle movement are calculated by tracking the movement of embedded fluorescent microspheres at each node (red points) of a user-defined grid superimposed on the image. (D) A displacement field can be calculated from the nodal displacements. Here, the series of images depicts the evolution of substrate displacement vectors, where the arrows indicate direction and the displacement magnitudes are color coded.

### Microneedle-Induced PA Gel Deformations

Controlled local mechanical deformations in the PA gel substrate were produced using a sterile 0.25 mm × 75mm acupuncture needle (Tai-Chi Brand, Suzhou Shenlong Medical Apparatus Co., Ltd., Suzhou, China) secured to a high-resolution linear actuator (M-230.25, PI (Physik Instrumente) LP, Auburn, MA) and inserted into the gel. The actuator was controlled with MTESTQuattro software (ADMET, Norwood, MA), and mounted to a high-precision, manual linear stage (462 XYZ-M ULTRAlign, Newport Corp., Irvine, CA).

Single keratinocytes that were at least 350 μm away from any neighboring cells were located and imaged. Circles measuring either 100 μm or 200 μm in radius and centered on the cell nucleus were superimposed on the live image using Nikon Elements software. The needle tip was then inserted to a depth of approximately 40 μm below the surface of the gel at a location on the circle that was intentionally irrespective of the direction of cell migration (Figure 1B). The needle was then displaced at a constant rate of 1 μm/min in order to mimic both the average instantaneous velocity of single keratinocytes and the rate of change of keratinocyte-induced substrate deformations we previously observed on soft PA gels [42].

Following needle insertion, time-lapse images were acquired at five-minute intervals for two hours, roughly the time at which time the needle began to exit the image field of view. As many as three sequential experiments were conducted on each gel, such that all observations were conducted within 10 hours of the start of the experiment. The two hour imaging period for any given cell was determined to be of acceptable duration because it was substantially larger than the average persistence time characteristic of a keratinocyte migrating on a soft PA gel (14.4 ± 17 minutes). To calculate this persistence time, a parallel set of control experiments were conducted where the needle was absent and images were acquired every five minutes over 24 hours. The mean squared displacements calculated from these time-lapse images were fit to the persistent random walk model of cell motility [53-55] using the method of Wu *et al*. [55] and averaged to extract the persistence time.

### Chemical Inhibitors

The Rho kinase inhibitor, Y27632 (ALX-270-333-M001, Enzo Life Sciences, Inc., Farmingdale, NY) [24-27, 56], and a NM II inhibitor, blebbistatin (ab120425, Abcam, Cambridge, MA) [57, 58], were added to the culture medium of some experiments in order to assess how inhibiting components of the mechanosensing and force generating machinery of the cytoskeleton affected keratinocyte behavior in response to the needle-induced substrate deformations. One hour after the cells were added to the PA gels (*i.e*., 2 hours before imaging began) either Y27632 or blebbistatin was added to the culture medium to produce a final concentration of 50 μM of inhibitor.

### Experimental Conditions

Four different experimental conditions involving needle-induced substrate deformations were investigated, where the initial distance of the needle away from the cell (Soft-100, Soft-200) and the effects of chemical inhibition (Soft-Y27632, Soft-Bleb) on cell migration were investigated (Table 1). An additional control condition (Soft-Control), where the needle was absent, was also examined. Ten cells were analyzed for each condition.

**Table 1.** Experimental Conditions

| Condition | Gel | Needle Location | Chemical Inhibitor | n |
| --- | --- | --- | --- | --- |
| Soft-100 | 1.2 kPa | 100 $\mu\text{m}$ | - | 10 |
| Soft-200 | 1.2 kPa | 200 $\mu\text{m}$ | - | 10 |
| Soft-Y27632 | 1.2 kPa | 100 $\mu\text{m}$ | Y27632 50 $\mu\text{M}$ | 10 |
| Soft-Bleb | 1.2 kPa | 100 $\mu\text{m}$ | Blebbistatin 50 $\mu\text{M}$ | 10 |
| Soft-Control | 1.2 kPa | - | - | 10 |

### PA Gel Substrate Displacement Tracking

PA gel substrate displacements were measured, as was done previously [42], by using a custom MATLAB (Mathworks, Inc, Natick, MA) algorithm based on spatial cross-correlation between subsequent image pairs of the embedded fluorescent microspheres [59]. Specifically, an array of evenly spaced grid points was placed on a subset of the image that contained the cell and extended to the location of microneedle insertion. Subwindows around each grid point were then correlated between image pairs to obtain the corresponding displacements in the substrate. The grid point positions were updated and used to generate the subwindows for correlation between the next image pair. In this manner, the grid point positions track with the microspheres in the deforming substrate. Thus, the 2D position vector of the substrate, *r̄*_*s*_(*t*), and the incremental substrate displacement between frames, *ū*_*s*_(*t*) = *r̄*_*s*_(*t*) – *r̄*_*s*_(*t* – 1), at time, *t*, are straightforward to describe.

### Cell Motility

The 2D position of the cell, *r̄*_*c*_(*t*), was determined by hand tracking the location of the nucleus in each image using the Manual Tracking plugin in ImageJ (National Institutes of Health, Bethesda, MD). In the experiments involving the needle, it was necessary to distinguish between cell movement from cell motility versus cell movement stemming from needle-induced deformations in the substrate. Thus, the total incremental displacement of the cell, *ū*_*c*_(*t*), between image pairs is given by:

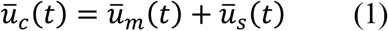

where *ū*_*m*_ is the incremental displacement vector of the cell due to cell motility and *ū*_*s*_ is the incremental displacement vector of the underlying substrate. Here, *ū*_*s*_ was made coincident with the position of the cell via linear interpolation from the displacements measured in the surrounding set of tracked grid points. For the limiting case of no substrate displacement (*i.e*., when there is no needle and *ū*_*s*_(*t*) = 0), cell movement is from cell motility alone. The position vector of the cell due to cell migration alone, *r̄*_*m*_, was then calculated by substituting the relationship *ū*_*m*_(*t*) = *r̄*_*m*_(*t*) – *r̄*_*m*_(*t* – 1) into Eq. (1) and rearranging to give:

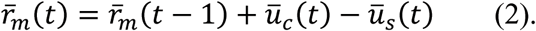

Both *r̄*_*m*_ and *ū*_*m*_, which we refer to as the adjusted position vector and adjusted cell displacement vector, respectively, were used in the data analysis described below.

### Directional Analysis and Circular Statistics

We posit that if a cell is preferentially migrating in a direction coincident with the direction of substrate displacement ‒ as opposed to changing direction randomly ‒ then the distribution of the direction vectors should deviate significantly from a uniform distribution (*i.e*., random cell migration). Furthermore, the distribution should have a mean direction that coincides with the mean direction of substrate displacement. To determine if such a relationship was present, an angle distribution of the direction of cell migration was obtained by calculating the angle of the adjusted cell displacement vector, *θ*_*ū*_*m*__, which is given by *θ*_*ū*_*m*__ = sin^‒1^(*u*_*m*_*y*__/*u*_*m*_*y*__), where *u*_*m*_*x*__, and *u*_*m*_*y*__ are the components of *ū*_*m*_. In a similar manner, the angle distribution of the direction of substrate displacement, *θ*_*ū*_*s*__, was calculated from *ū*_*s*_. Both *θ*_*ū*_*s*__ and *θ*_*ū*_*m*__ were calculated for each frame, binned together for all ten experiments associated with a given condition, and represented as polar histograms. The CircStat tool box for MATLAB [60] was then used to compare the two distribution via a modified Rayleigh test for uniformity, or V test [61], to determine if the distribution for *θ*_*ū*_*m*__ was random (*i.e*., distributed uniformly around a circle), or if the distribution for *θ*_*ū*_*m*__ was not random and its mean direction coincided with the average direction of *θ*_*ū*_*s*__.

In addition, we also calculated the alignment angle, *ϕ*, between *ū*_*m*_ and *ū*_*s*_ [62]. Here, *ϕ* ranges between *ϕ* = 0° and *ϕ* = 180°, which corresponds to prefect alignment between *ū*_*m*_ and *ū*_*s*_ in the same or opposite directions, respectively. The cumulative probability distribution function of the alignment angle, *P̄*(*ϕ*), was then constructed. Here, if *ϕ* is uniformly distributed, meaning that there is no preferential alignment toward the direction of underlying substrate, all angle are equally likely and the probability function will be a straight line with probability 0 at 0° to probability 1 at 180°. If *P̄*(*ϕ*) increases faster than the case of a straight line, it means that the cells are biased towards the direction of substrate deformations.

## RESULTS

Control experiments without the insertion of a needle into the gel were performed in order to obtain baseline information on single keratinocyte migration behaviors. These cells dynamically interrogated and deformed the substrate as they moved randomly all over the gel surface (Movie 1). The average cell speed, net distance traveled, and maximum distance traveled over the 24 hour period was 1.27 ± 0.25 μm/min, 178.14 ± 123.68 μm, and 238.16 ± 115.05 μm, respectively (Table 2). The persistence time of these cells was calculated as 14.4 ± 17.0 minutes. Consequently, a duration of two hours for each microneedle experiment was deemed more than sufficient for determining whether directed migration towards the needle was occurring for each condition examined.

**Table 2.**
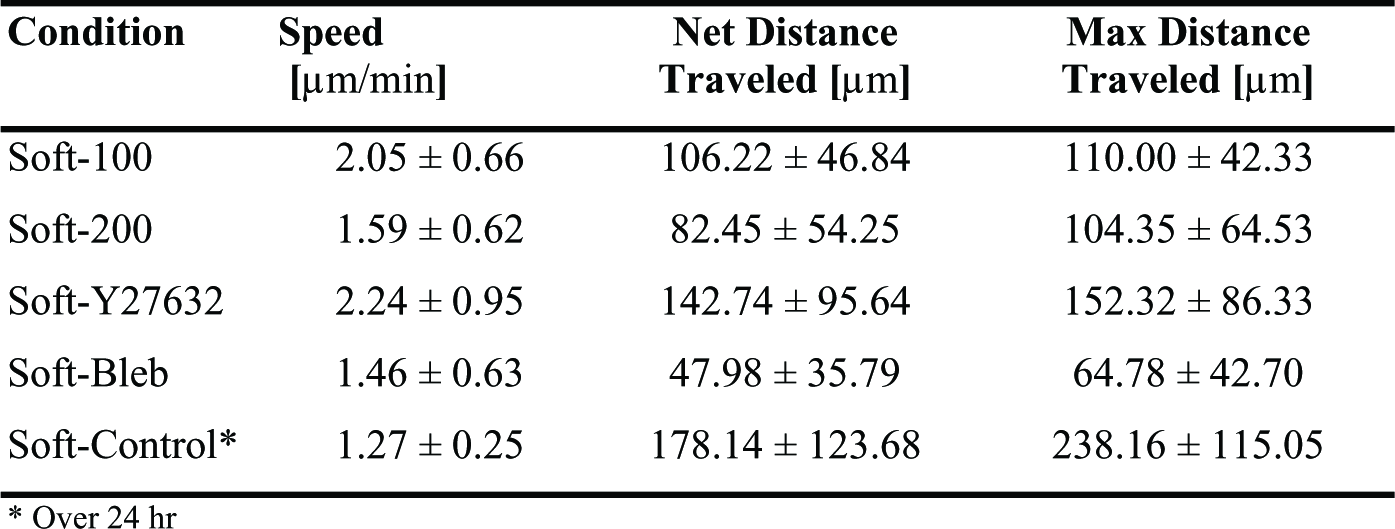
Baseline Keratinocyte Migration Metrics on a Non-Deformed Substrate Results

| Condition | Speed<br>[$\mu\text{m}/\text{min}$] | Net Distance<br>Traveled [$\mu\text{m}$] | Max Distance<br>Traveled [$\mu\text{m}$] |
| --- | --- | --- | --- |
| Soft-100 | $2.05 \pm 0.66$ | $106.22 \pm 46.84$ | $110.00 \pm 42.33$ |
| Soft-200 | $1.59 \pm 0.62$ | $82.45 \pm 54.25$ | $104.35 \pm 64.53$ |
| Soft-Y27632 | $2.24 \pm 0.95$ | $142.74 \pm 95.64$ | $152.32 \pm 86.33$ |
| Soft-Bleb | $1.46 \pm 0.63$ | $47.98 \pm 35.79$ | $64.78 \pm 42.70$ |
| Soft-Control* | $1.27 \pm 0.25$ | $178.14 \pm 123.68$ | $238.16 \pm 115.05$ |

\* Over 24 hr

Next, the servo-controlled needle was inserted into the gel a nominal distance of 100 μm away from the cell nucleus and displaced at a constant rate of 1 μm/min to produce a constant and repeatable deformation in the gel over a 2 hour period (Movie 2). These cells, which prior to the insertion of the needle were migrating in random directions, typically altered course and moved in the direction of needle displacement (Fig. 2). This behavior was also observed, though to a lesser extent, in experiments where the needle was nominally inserted 200 μm from the cell center (Movie 3).

**Figure 2.**
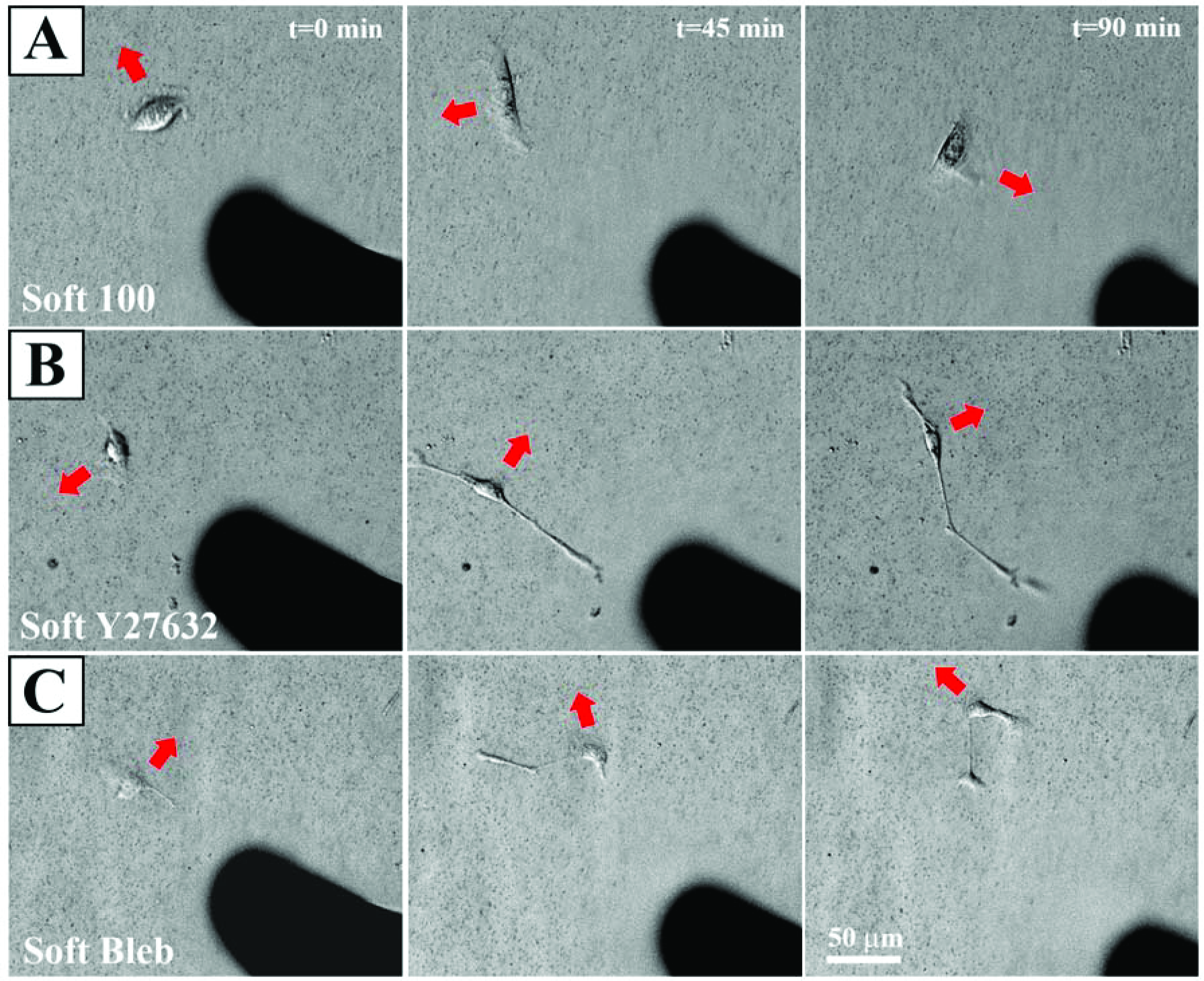
Direction of keratinocyte migration in response to needle-induced substrate deformations at t = 0 min, t = 45 min, and t = 90 min. Red arrows indicate the current direction of cell movement. (A) For Soft 100, the cell, which is initially migrating away from the needle gradually reorients and moves in the direction in which the needle is pulling the substrate. (B) For Soft Y27632 and (C) Soft Bleb, the cells move in a manner that is unresponsive to the substrate deformations produced by needle movement. Also, note that exposure to both drugs perceptibly altered cell morphology.

The addition of Y27632 (Movie 4) and blebbistatin (Movie 5) altered keratinocyte morphology and migration behavior (Fig. 2). Both drugs induced the formation of highly extended cells with long, thin processes, but their respective migration patterns differed appreciably. Most importantly, keratinocytes in both cases were unresponsive to substrate deformations produced by the needle.

Post-experiment analysis found no significant differences in the magnitude or rate of substrate displacements induced by the needle amongst the four experimental conditions. However, the true needle tip to cell nuclei distance for circles measuring 100 μm and 200 μm was less than the nominal distance at 84.0 μm ± 5.7 μm, 85.9 μm ± 4.4 μm, 86.4 μm ± 6.0 μm, and 182.3 μm ± 5.4 μm for Soft-100, Soft-Y27632, Soft-Bleb, and Soft-200, respectively.

In order to confirm that the keratinocytes were migrating towards the needle and not simply carried by the underlying deforming substrate, we calculated the adjusted cell position and plotted the adjusted cell paths for all experiments (Fig. 3). These paths clearly show that the cells primarily migrated in the direction of the needle and that exposure to either Y27632 or blebbistatin blocked this effect. A comparison of the distribution of *θ*_*ū*_*m*__ to the direction of needle displacement (Fig. 4) also quantitatively and statistically supported this notion. V tests confirmed that for both Soft-100 and Soft-200 the direction of cell movement coincided with the direction of needle movement (p < 0.001). V tests also indicated that the direction of cell movement for Soft-Y27632 and Soft-Bleb samples was not significantly different from a uniform distribution and did not coincide with the direction of needle-induced substrate displacements.

**Figure 3.**
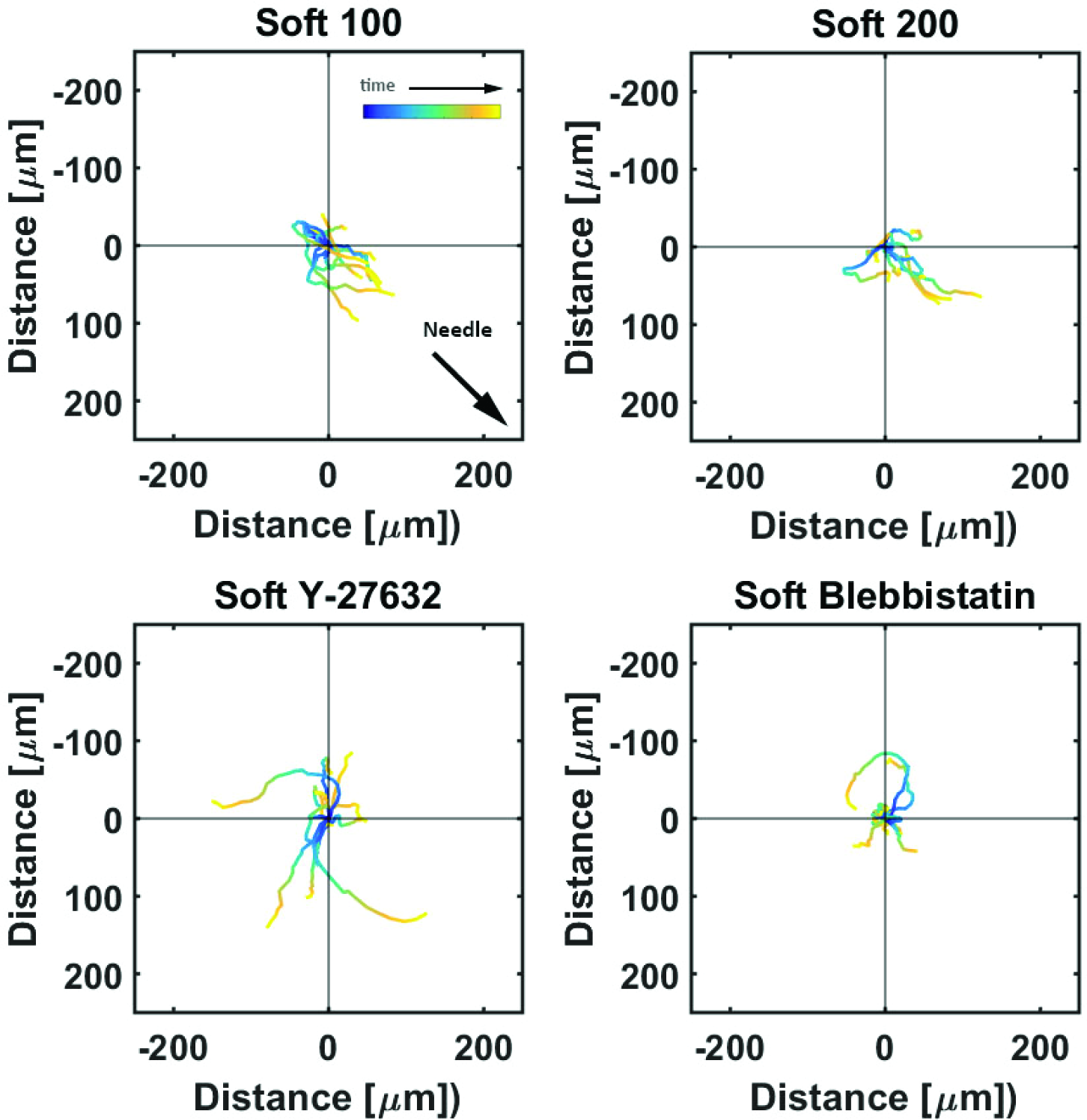
Adjusted Cell Motility on PA gels. Each color line shows the trace of ***r̄***_***m***_, the position vector of the cell due to cell migration effects alone (see Eq. 2). Each cell path is color-coded to show the relative time along the path from its beginning at the origin (dark blue) to the final tracked position (yellow). The majority of cells moved toward the displacing needle for needles nominally positioned either 100 μm or 200 μm from the cell centroid at the start of the experiment (i.e., Soft 100 and Soft 200, respectively). Exposure to the rho kinase inhibitor Y27632 or the actin-myosin inhibitor blebbistatin interrupted this directed migration.

**Figure 4.**
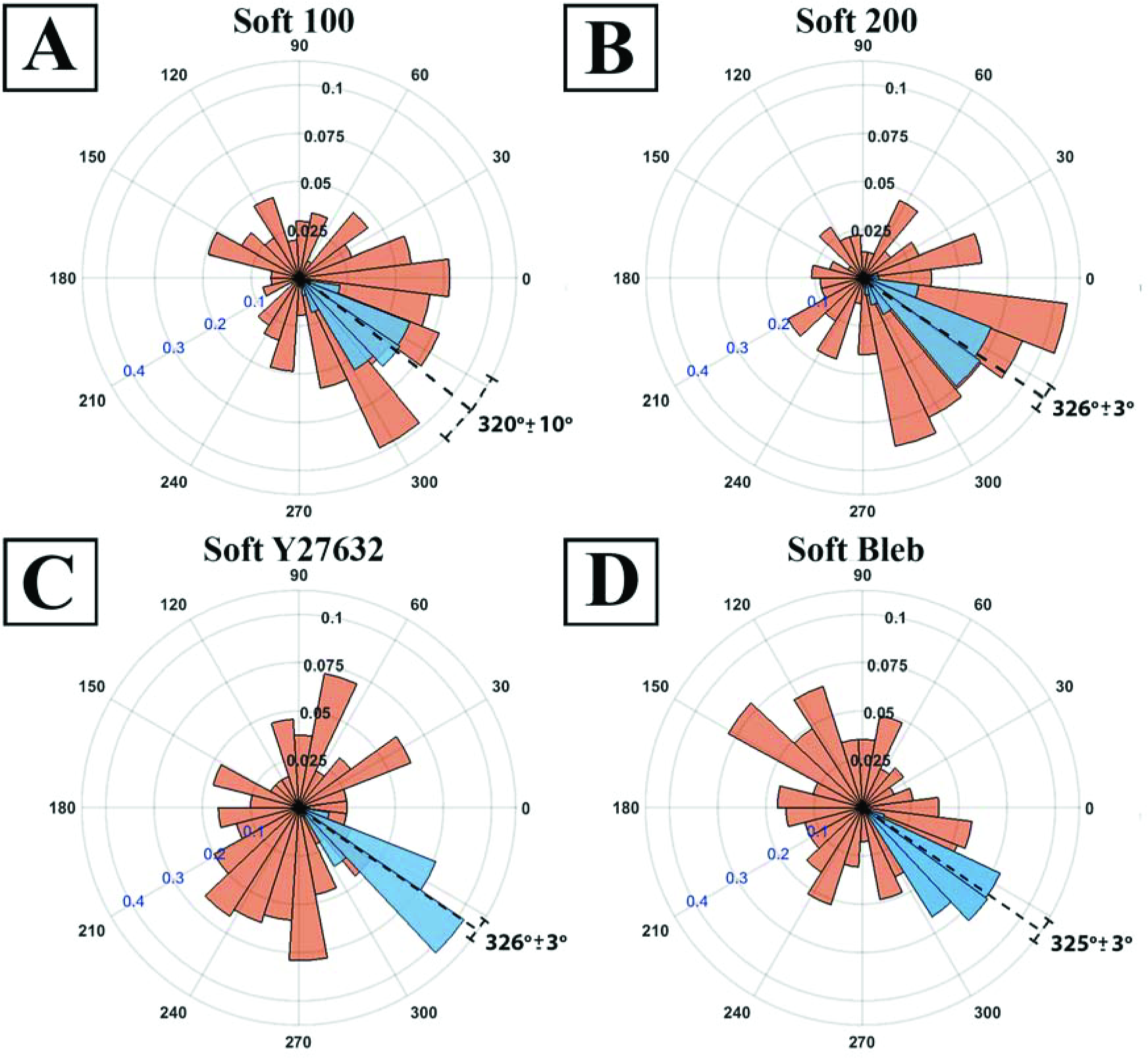
Polar histograms showing the combined angle distributions in degrees of adjusted cell displacement ***θ***_***ū***_***m***__ (red) and substrate displacement ***θ***_***ū***_***s***__ (blue) for all 10 replicates of the (A) Soft-100, (B) Soft-200, (C) Soft-Y27632, and (D) Soft-Bleb experiments. Each distribution is normalized by the total number of observations with the black radial text associated with ***θ***_***ū***_***m***__ and the blue radial text associated ***θ***_***ū***_***s***__. The dashed line shows the mean and standard deviation of the angle of needle movement. For all four conditions, the displacements of the substrate aligned with the direction of needle movement. In addition, for Soft-100 and Soft-200, V-tests indicate that the direction of cell movement is coincident with that of the needle-induced substrate deformations, but not for soft-Y27632 or Soft-Bleb (p <0.001).

**Figure 5.**
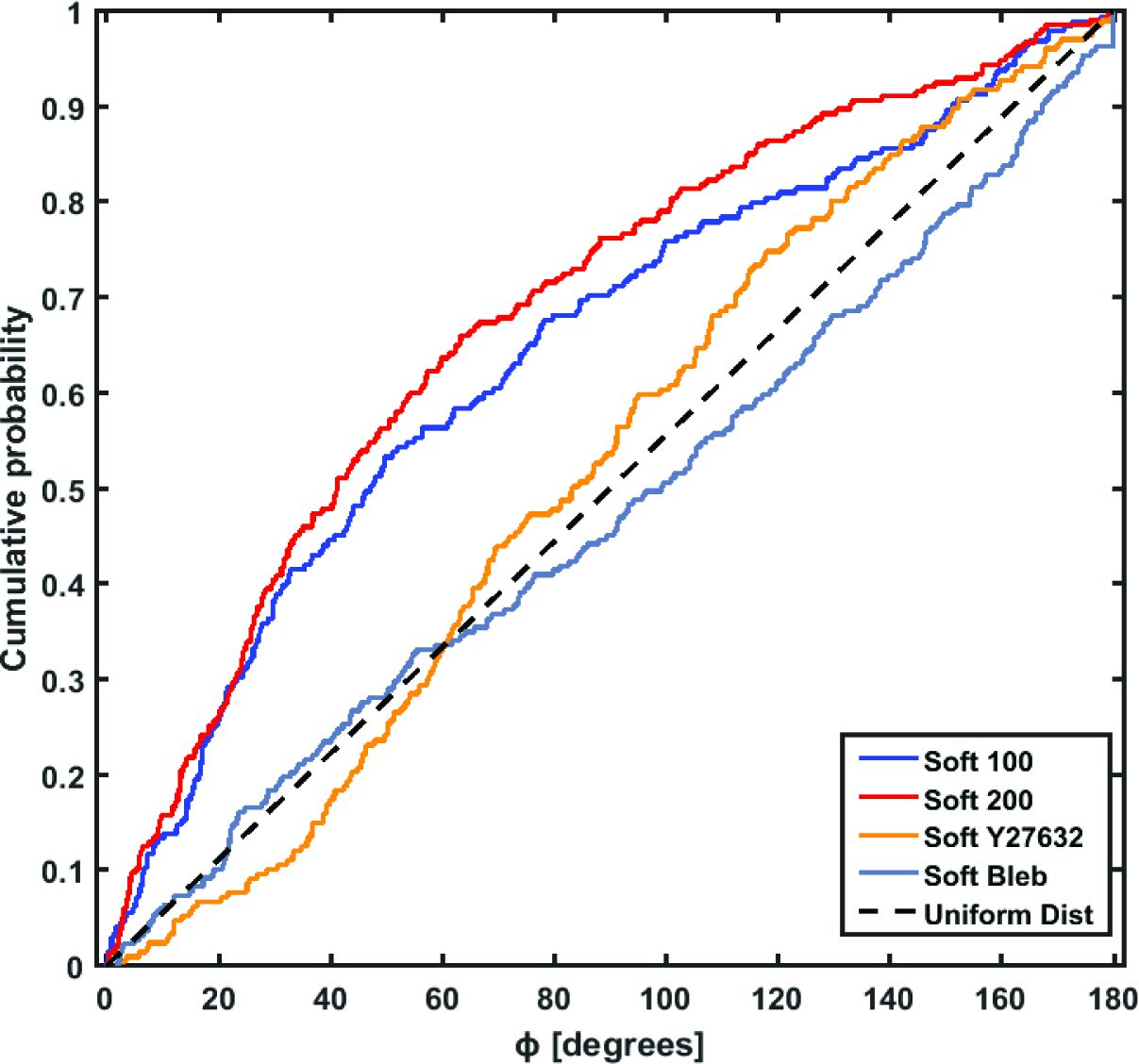
The cumulative probability distribution (CPD) of ***ϕ***, the angle between *θ*_***ū***_***m***__ and *θ*_***ū***_***s***__. Random cell migration results in a uniform angle distribution where the CPD traces out a straight line (dotted line). The higher probabilities for low values of ***ϕ*** for Soft-100 (blue) and Soft-200 (red) indicate directed movement of the cells towards the direction of needle-induced substrate deformations. The CPD for Soft-Y27632 (orange) and Soft-Bleb (gray) more closely follow that of a uniform distribution.

A third piece of evidence supporting the idea that keratinocytes respond to mechanical cues in the substrate comes from an examination of *P̄*(*ϕ*) (Fig. 4). *P̄*(*ϕ*) for both Soft-100 and Soft-200 increased faster than the straight line representative of a uniform distribution, indicating that the cells were biased towards the direction of needle-induced substrate displacements. In contrast, *P̄*(*ϕ*) for Soft-Y7632 and Soft-Bleb more closely approximated a uniform distribution, indicating that these inhibitors negated the influence of mechanical cues on keratinocyte migration.

## DISCUSSION

During wound healing, the composition and structure of the wound bed transitions dramatically from that of the initial provisional fibrin clot to a dense, vascularized, cellular bed of granulation tissue, to a final state of remodeled replacement ECM [63]. Concurrently, the mechanical properties of the wound site change continuously, with atomic force microscope (AFM) measurements on wounded rat skin showing increases in stiffness from 18.5 kPa at day 7 to 29.4 kPa by day 9 [64], compared to median measurements in human papillary and reticular dermis of 0.82 and 1.12 kPa, respectively [65]. Given that a variety of mechanical cues and physical contexts haven been shown to regulate physiological and pathophysiological cellular behaviors [48, 66-70], it is reasonable to ask whether such cues also have the potential to guide temporal and spatial aspects of keratinocyte re-epithelialization of the wound site.

Here, we set out to validate the hypothesis that keratinocytes adherent to soft substrates can sense and respond to substrate deformations, and that this is the explanation for the enhanced cooperativity in de novo epithelial sheet formation that we observed in our previous work [42]. Towards this end, we designed an experiment in which we experimentally imposed substrate deformations similar in magnitude and rate of deformation to those that can be generated by a single keratinocyte or evolving multicellular aggregate of keratinocytes. By imposing these experimental deformations within a defined length scale relative to an otherwise randomly migrating isolated keratinocyte, we were able to consistently induce substrate deformation-directed keratinocyte migration. Furthermore, using a Rho kinase inhibitor (Y27632) and a NM II inhibitor (blebbistatin), we have demonstrated that the Rho/ROCK pathway and the engagement of NM II are both essential to the observed process of substrate deformation-directed keratinocyte migration.

Our observations are consistent with the pioneering work of Lo *et al*., who, in addition to showing that cells preferentially migrate up gradients of rigidity (*i.e*., durotaxis), also used a blunted microneedle to demonstrate that localized mechanical deformations in the substrate could direct 3T3 fibroblast migration either towards or away from the needle, depending on whether the substrate was pulled or pushed, respectively [70]. More recently, Plotnikov *et al*. obtained similar results following the same methodology, where the needle was inserted 10 μm away from the cell edge, and then instantaneously stretched to the elastic limit and held in place for 45 minutes [45]. In contrast, we applied a controlled, constant rate of substrate deformation with the needle inserted 100 μm and 200 μm from the cell center and quantitatively tracked both the imposed substrate deformations and the cell migration cell path over a 2 hour period. Although the cellular response was essentially the same, the nature of the mechanical cues involved was temporally and spatially different from these earlier studies. Because the microneedle imposed substrate deformations at a constant rate, and cells were migrating in random directions with respect to the subsequently imposed substrate deformation, the potential mechanical cues a given cell experienced were varied. The diversity of spatial and temporal mechanical inputs likely contributed to the variability in the cell paths taken (Fig. 3), where not all followed the needle-induced deformations for the duration of the experiment.

The mechanism(s) underlying the aggregation of single cells on deformable substrates is likely to be similar, if not identical, to the mechanisms that gave rise to the substrate deformation-induced keratinocyte migration patterns observed in our experiments. Multicellular aggregating behavior on soft but not stiff PA gels is a phenomenon that has been observed by others [43, 44]. In particular, Reinhart-King *et al*. [43] found that endothelial cell pairs on soft PA gels were able to interact with each other by sensing mechanical forces exerted between them through the substrate. The dispersion, or tendency of these cell pairs to migrate away from their initial location, was reduced significantly compared to that of single cells. Guo *et al*. posited that this type of behavior could be explained in terms of comparisons between mechanical signals from the cell and the substrate and mechanical signals between the cell and neighboring cells [44]. Cells migrate away from each other if the mechanical signals are greater in the substrate than from neighbors, and towards each other when the signals are reversed. The tendency for keratinocytes to migrate in the direction of needle displacement observed here is consistent with this hypothesis.

A common explanation for what guides directed cell migration is that deformation of the substrate either by a needle or an adjacent cell or aggregate of cells produces a durotactic gradient in gel rigidity via nonlinear strain stiffening. PA gels, however, are generally considered linear elastic materials, which implies that local gel rigidity does not change with increasing strain. It is possible that the deformations in these studies were large enough to exceed the linear elastic range of the gel, such that strain stiffening did occur, especially, when one considers that nonlinear strain stiffening has been reported for deformations of as little as 2-6 μm [71]. At 70 μm, gel displacements in our study were approximately an order of magnitude higher, which suggests that strain stiffening may have occurred. Whether the cells are responding to a gradient in substrate stiffness, or some other mechanical/environmental cue, however, remains unclear. With respect to the latter, it is also possible that substrate deformations produced a gradient in potential substrate adhesive contacts (*i.e*., haptotaxis) that resulted from increases in ECM protein density within areas of increasing substrate deformation. Alternatively, changes in gel thickness could also alter the gel topography in a way that favors directed cell migration towards the needle. We did not explicitly check for either of these possibilities, but plan to do so in future investigations, where we will also investigate the effect of varying the deformation rate and deformation profile (e.g., cyclical) on the keratinocyte’s migratory response.

Regardless of which mechanical cue(s) are directly responsible, the addition of either Y27632 or blebbistatin interrupted directed cell migration towards the needle-induced substrate deformation, supporting the notion that myosin contraction at anchoring junctions is a vital component of the mechanosensing apparatus. Similarly, exposure to both of these drugs also prevented multicellular aggregate formation on soft PA gels [44]. Several studies have shown that intracellular tension via actin-myosin engagement is necessary for the integration of key mechanosensing proteins (e.g., vinculin) into focal adhesions [13, 16, 72], and that interrupting myosin II contractility via exposure to blebbistatin or Y27632 interferes with this process [16, 45].

## CONCLUSION

In this work, we investigated the keratinocyte’s migratory response to controlled substrate deformations originating ~100 μm and ~200 μm away from an otherwise randomly migrating cell. Most keratinocytes altered their migration pathway, preferring a course that vectorially aligned with the direction of substrate deformation. Addition of either Y27632 or blebbistatin impaired the cellular migratory response to these mechanical deformations. Collectively, these results provide strong evidence supporting the notion that keratinocytes are mechanosensitive cells, further implying that the mechanobiology of keratinocytes could be an important factor that contributes to the process of re-epithelialization in states of both health and disease.

## ACKNOWLEDGEMENTS

Support for this work was provided by the National Science Foundation (CAREER 1452728) to E.A.S. In addition, J.C.S. acknowledges the Dermatology Foundation for their support of this work through a career development award.

